# Tripolar mitosis drives the association between maternal genotypes of *PLK4* and aneuploidy in human preimplantation embryos

**DOI:** 10.1101/182303

**Authors:** Rajiv C. McCoy, Louise J. Newnham, Christian S. Ottolini, Eva R. Hoffmann, Katerina Chatzimeletiou, Omar E. Cornejo, Qiansheng Zhan, Nikica Zaninovic, Zev Rosenwaks, Dmitri A. Petrov, Zachary P. Demko, Styrmir Sigurjonsson, Alan H. Handyside

## Abstract

Aneuploidy is prevalent in human preimplantation embryos and is the leading cause of pregnancy loss. Many aneuploidies arise during oogenesis, increasing in frequency with maternal age. Superimposed on these meiotic aneuploidies are a range of errors occurring during early mitotic divisions of the embryo, contributing to widespread chromosomal mosaicism. Here we reanalyzed a published dataset comprising preimplantation genetic testing for aneuploidy in 24,653 blastomere biopsies from day-3 cleavage-stage embryos, as well as 17,051 trophectoderm biopsies from day-5 blastocysts. We focused on complex abnormalities that affected multiple chromosomes simultaneously, seeking to quantify their incidences and gain insight into their mechanisms of formation. In addition to well-described patterns such as triploidy and haploidy, we identified 4.7% of day-3 blastomeres possessing karyotypes suggestive of tripolar mitosis in normally-fertilized diploid zygotes or descendant diploid cells. We further supported this hypothesis using time-lapse data from an intersecting set of 77 cleavage-stage embryos. The diploid tripolar signature was rare among day-5 blastocyst biopsies (0.5%), suggesting that complex aneuploidy generated by tripolar mitosis impairs cellular and/or early embryonic survival. Strikingly, we found that the tripolar mitosis mechanism is responsible for the previously described association with common maternal genetic variants spanning *PLK4*. Our findings are consistent with the role of *PLK4* as the master regulator of centriole duplication with a known capacity to induce tripolar mitosis when mutated or mis-expressed. Taken together, we propose that tripolar mitosis is a key mechanism generating karyotype-wide aneuploidy in cleavage-stage embryos and implicate *PLK4*-mediated centrosome abnormality as a factor influencing its occurrence.

## Introduction

Aneuploidy is common in human preimplantation embryos and increases in frequency with maternal age (1). While many aneuploidies originate in meiosis—primarily in females (oogenesis) rather than males (spermatogenesis) (2,3)—a substantial proportion arise after fertilization (postzygotic) due to errant chromosome segregation during early cleavage divisions (4). These mitotic errors produce mosaic embryos with two or more cell lineages possessing distinct chromosomal complements. Because aneuploidy is associated with negative pregnancy outcomes, many patients undergoing *in vitro* fertilization (IVF) treatment for infertility have their embryos tested by copy number analysis of all 24 chromosomes. Testing is applied to single or small numbers of biopsied cells with the aim of transferring only embryos that are euploid—an approach previously known as preimplantation genetic screening (PGS) or preimplantation genetic diagnosis for aneuploidy (PGD-A) but now termed preimplantation genetic testing for aneuploidy (PGT-A) (5,6).

Despite substantial improvements in detecting aneuploidy at the single cell level, distinguishing aneuploidies of meiotic and mitotic origin remains challenging because the chromosomal signatures can be similar. In some cases, discrimination may be achieved by incorporating parental genotype information to determine the parental origin of each embryonic chromosome. Specifically, observation of both maternal (or, rarely, paternal) haplotypes transmitted in a homologous region of an embryonic chromosome—a signature we term ‘both parental homologs’ or ‘BPH’—provides strong evidence of the meiotic origin of a trisomy (Fig. 1A). Meanwhile, because aneuploidy is rare in sperm (1-4%) (7) and paternal BPH affects only 1% of preimplantation embryos (8), aneuploidies involving gain or loss of paternal homologs are predominantly mitotic errors. To date, several methods have been developed to infer the transmission of individual parental homologs based on single nucleotide polymorphism (SNP) microarray data, facilitating classification of meiotic and mitotic aneuploidies. These include Parental Support (9,10) and karyomapping (11,12), which have recently been validated for linkage-based preimplantation genetic diagnosis (PGD) of single gene disorders (13,14).

**Figure 1.**
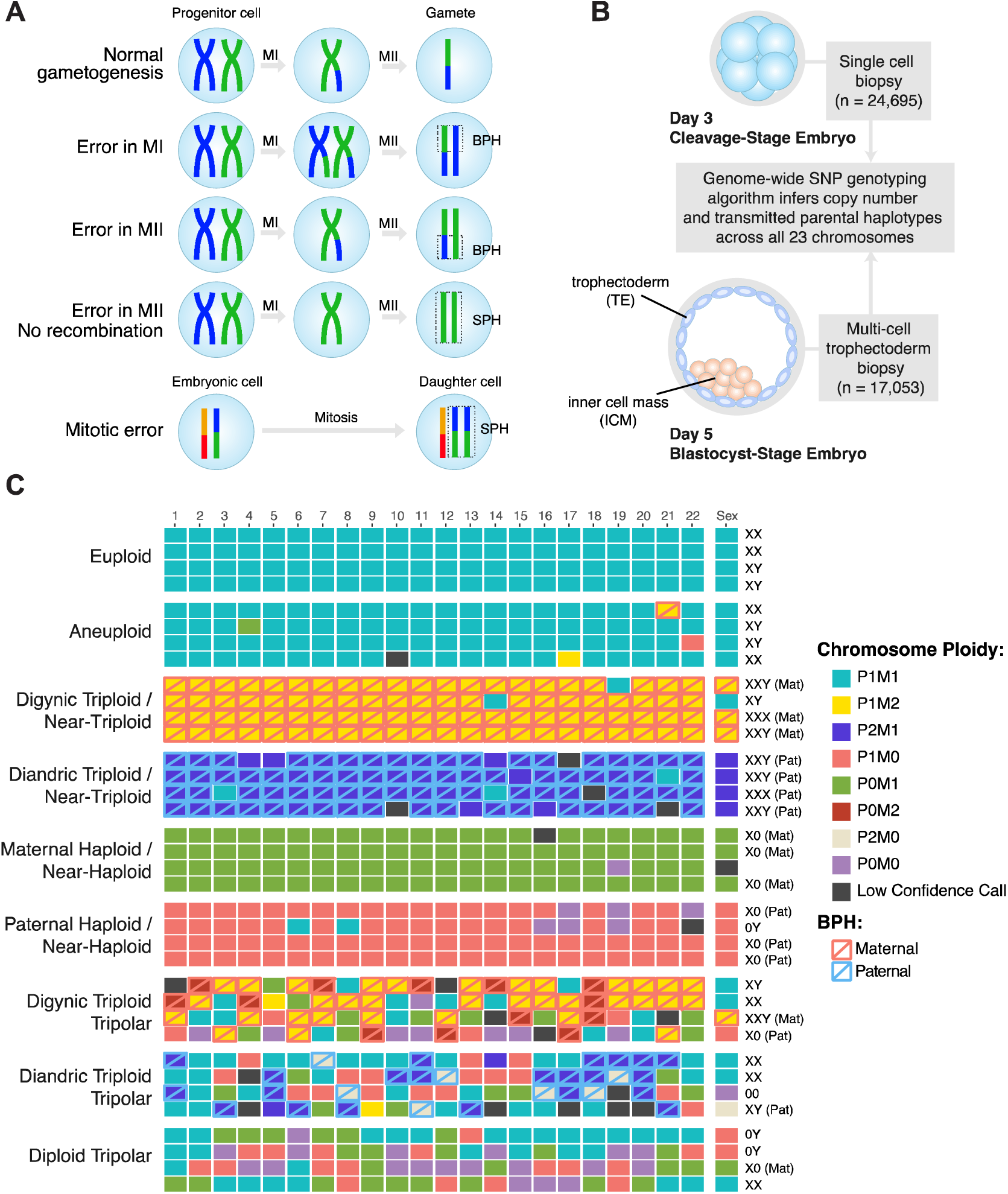
Patterns of karyotype-wide chromosome abnormalities observed in the preimplantation human embryos. (A) Schematic describing the signature of meiotic chromosome gain, adapted from Rabinowitz et al. (10) and McCoy et al. (8). The detection of non-identical homologous chromosomes inherited from a single parent (a signature that we term ‘both parental homologs’ or ‘BPH’) indicates a meiosis I (MI) or meiosis II (MII) error. Chromosome gains that involve identical chromosomes from a single parent (termed ‘single parental homolog’ or ‘SPH’) can occur either due to mitotic errors or MII errors in the absence of recombination. (B) Schematic cross-section showing the sample sizes of cleavage and blastocyst-stage embryo biopsies which underwent genome-wide pre-implantation genetic screening. Both parents were also genotyped using the same single nucleotide polymorphisms (SNP) microarray platform to facilitate inference of chromosome copy number and determine the parental origin of each chromosome. This approach also enables classification of trisomies of meiotic origin. (C) Examples of patterns of whole-chromosome copy number variation observed in the dataset. Most samples were normal (euploid) or contained a few single-chromosome imbalances (aneuploid). However, complex karyotype-wide abnormalities were also relatively prevalent (12.0%; Table 1). Each column represents a chromosome (1-22; left to right along with the sex chromosomes at the far right). Each row represents a distinct embryo biopsy. Four representative biopsies are depicted for each category of chromosome abnormality. M: maternal, P: paternal, BPH: both parental homologs.

**Table 1.** Incidence of aneuploidy and karyotype-wide abnormality in day-3 (d3) blastomere biopsies and day-5 (d5) trophectoderm biopsies.

|  | Day-3 blastomere biopsies |  |  |  | Day-5 trophectoderm biopsies |  |  |  |  |  |
| --- | --- | --- | --- | --- | --- | --- | --- | --- | --- | --- |
| Category | N samples (%) | Meiot. (%)<sup>a</sup> | Mitot. (%)<sup>b</sup> | Ambig. (%)<sup>c</sup> | N samples (%) | Meiot. (%)<sup>a</sup> | Mitot. (%)<sup>b</sup> | Ambig. (%)<sup>c</sup> | (% d3) / (% d5) | Selection criteria |
| Euploid | 8188 (33.2%) |  |  |  | 8687 (50.9%) |  |  |  | 0.65 | All 23 chromosomes biparental disomic |
| Euploid/NA | 1173 (4.8%) |  |  |  | 774 (4.5%) |  |  |  | 1.05 | Biparental disomic with ≥1 low-confidence calls |
| Aneuploid | 11,167 (45.3%) | 3585 (14.5%) | 3116 (12.6%) | 4466 (18.1%) | 6707 (39.3%) | 2263 (13.3%) | 1419 (8.3%) | 3025 (17.7%) | 1.15 | ≥1 and <10 aneuploid chromosomes |
| Karyotype-wide abnormality | 4125 (16.7%) | 1706 (6.9%) | 2152 (8.7%) | 267 (1.1%) | 883 (5.2%) | 483 (2.8%) | 299 (1.8%) | 101 (0.6%) | 3.23 | ≥10 aneuploid chromosomes |
| Total<sup>d</sup> | 24653 | 5291 (21.5%) | 5268 (21.4%) | 4733 (19.2%) | 17051 | 2746 (16.1%) | 1718 (10.1%) | 3126 (18.3%) |  |  |

d3: single blastomere biopsy from day-3 cleavage-stage embryo; d5: multiple cell trophectoderm biopsy from day-5 blastocyst; NA: missing data, indicating a low-confidence call; <sup>a</sup>Samples possessing a putative meiotic error, defined as one or more chromosomes

These technical improvements in PGT-A methodologies have provided substantial insights into the mechanisms of aneuploidy formation. An adapted form of karyomapping, for example, recently revealed a novel mechanism of meiotic error termed ‘reverse segregation’, whereby sister chromatids separate at meiosis I, followed by (occasionally errant) segregation of non-sister chromatids at meiosis II (12). This adds to premature separation of sister chromatids (PSSC) and meiotic non-disjunction as the predominant known mechanisms of meiotic error (12,15). Mitotic aneuploidies, meanwhile, have traditionally been attributed to mitotic non-disjunction, whereby chromatids fail to separate, as well as anaphase lag, whereby chromatids are lost after delayed migration toward the spindle poles (16).

A distinct class of severe mitotic aneuploidy, the origins of which have remained poorly understood, was previously referred to as ‘chaotic mosaicism’ due to its seemingly random chromosomal constitution (17). Though prevalent at the cleavage stage, chaotic mosaic embryos rarely achieve blastocyst formation (18). One hypothesized source of chaotic aneuploidy is the formation of multipolar mitotic spindles, which may cause the cell to cleave into three or more daughter cells. This phenomenon was recently confirmed by time-lapse analysis and karyomapping of 32 embryos, including cells biopsied from 25 arrested cleavage-stage embryos as well as cells extruded from seven additional developing embryos at the time of blastocyst formation (19). Strikingly, 90% of these cells exhibited evidence of chromosome loss (19), indicating that multipolar mitosis may constitute the primary mechanism inhibiting *in vitro* preimplantation development. Recognized since the foundational work of Boveri in the late 19^th^ century (20), multipolar spindles are in turn thought to form due to abnormalities of the centrosome. As the microtubule-organizing center (MTOC) that coordinates chromosome segregation, the centrosome requires tight regulation of its replication to ensure mitotic fidelity. Supernumerary (>2) centrosomes can lead to the formation of a multipolar rather than normal bipolar spindle (21). If proceeding through anaphase, replicated chromosomes may segregate among multiple daughter cells.

Excess centrosomes and tripolar mitosis are sometimes caused by abnormal fertilization. This includes fertilization with two sperm or retention of the second polar body, producing diandric or digynic tripronuclear (3PN) zygotes, respectively (22,23). However, tripolar spindles have also been observed by laser confocal microscopy both at the cleavage and blastocyst stages in embryos derived from normally-fertilized dipronuclear (2PN) zygotes (18), albeit at a lower rate than for 3PN zygotes. Indeed, tripolar mitosis (also called ‘trichotomic mitosis’ or ‘direct unequal cleavage [DUC]’) has recently been documented in a substantial proportion (17%–26%) of cleaving embryos based on time-lapse imaging (24,25,26,27), even after intracytoplasmic sperm injection (ICSI) which virtually eliminates polyspermic fertilization. Rather than abnormal centrosome transmission, these observations suggest possible dysregulation of centrosome duplication.

Here we leverage a published PGT-A dataset (28,8) to gain further insight into the molecular origins of chaotic aneuploidy. These data comprise 46,439 embryo biopsies genotyped by SNP microarray with aneuploidies inferred using the Parental Support algorithm (9) (Fig. 1B). This approach provides several advantages for the study of aneuploidy. The first advantage is the resolution of the data, which include estimates of copy number across all 24 chromosomes, as well as identification of meiotic chromosome gains. Second, the large sample size facilitates detection and quantification of rare forms of chromosome abnormality. Finally, by including both cleavage- and blastocyst-stage embryos, the dataset provides valuable information about the early developmental implications of various forms of aneuploidy. Based on our analysis, we provide evidence that tripolar mitosis of normally fertilized 2PN zygotes or their descendant cells is a key mechanism contributing to chaotic aneuploidy in cleavage-stage embryos. Furthermore, we demonstrate that this mechanism drives the association we previously reported (28) between mitotic-origin aneuploidy and common maternal genetic variants spanning *PLK4*, the master regulator of centrosome duplication.

## Results

### Mitotic-origin aneuploidy is prevalent in preimplantation human embryos

Excluding replicate and uninformative samples (≥10 chromosomes with low-confidence or nullisomic calls) we quantified various forms of chromosomal abnormality in 41,704 embryo biopsies from a total of 6319 PGT-A cases. Maternal age, including patients and egg donors, ranged from 18 to 48 (median = 37), while paternal age ranged from 21 to 77 (median = 39). Indications for PGT-A were diverse and included advanced maternal age, recurrent pregnancy loss, sex selection, previous IVF failure, male factor infertility, unexplained infertility, previous aneuploidy, and translocation carrier status (8).

The overall rate of euploidy among day-3 blastomere biopsies was 38.0% compared to 55.5% for day-5 trophectoderm biopsies (Table 1). We note that in light of mosaicism, which is common during preimplantation development (1,4,29) ploidy status of the embryo biopsy may not necessarily reflect the ploidy of the rest of the embryo. Among samples with 1-3 aneuploid chromosomes, a total of 2579 (30.2%) day-3 blastomere samples possessed at least one putative meiotic (BPH) chromosome gain, compared to 1915 (32.7%) day-5 trophectoderm samples. Conversely, a total of 1880 (22.0%) day-3 blastomere samples and 1069 (18.2%) day-5 trophectoderm samples possessed only putative mitotic errors (loss or SPH gain of ≥1 paternal chromosome with no co-occurring maternal or paternal BPH gains).

### Karyotype-wide abnormalities are more common in cleavage-than blastocyst-stage embryos

In addition to these less severe aneuploidies, we sought to investigate the origin of ‘karyotype-wide’ abnormalities that affect many chromosomes simultaneously. These include so-called ‘chaotic’ aneuploidies, characterized by seemingly random chromosome complements (17). We identified 5008 (12.0%) embryo samples exhibiting patterns of karyotype-wide chromosome abnormalities (arbitrarily defined as ≥10 aneuploid chromosomes), a subset of which could be classified into distinct categories (Fig. 1C; Table 2). The proportion of these karyotype-wide abnormalities was significantly greater in day-3 blastomere samples (16.7%) than in day-5 trophectoderm samples (5.2%; Chi-Squared Test: ?2[1, *N* = 41,704] = 1272, *P* < 1 × 10^-10^).

**Table 2.** Frequencies of various forms of karyotype-wide abnormality in day-3 (d3) blastomere biopsies and day-5 (d5) trophectoderm biopsies.

| Type of karyotype-wide abnormality | No. d3 samples (%) | No. d5 samples (%) |
| --- | --- | --- |
| Digynic Triploid / Near-Triploid | 740 (3.0%) | 321 (1.9%) |
| Diandric Triploid / Near-Triploid | 25 (0.1%) | 32 (0.2%) |
| Maternal Haploid / Near-Haploid | 288 (1.2%) | 84 (0.5%) |
| Paternal Haploid / Near-Haploid | 126 (0.5%) | 25 (0.1%) |
| Multiple Maternal Gains (Non-Triploid) | 143 (0.6%) | 55 (0.3%) |
| Multiple Paternal Gains (Non-Triploid) | 16 (0.1%) | 4 (0.0%) |
| Both Maternal and Paternal Gains (Non-Triploid) | 61 (0.2%) | 38 (0.2%) |
| Multiple Maternal Losses (Non-Haploid) | 98 (0.4%) | 19 (0.1%) |
| Multiple Paternal Losses (Non-Haploid) | 79 (0.3%) | 11 (0.1%) |
| Both Maternal and Paternal Losses (Non-Haploid) | 137 (0.6%) | 5 (0.0%) |
| Digynic Triploid Tripolar | 93 (0.4%) | 13 (0.1%) |
| Diandric Triploid Tripolar | 25 (0.1%) | 0 (0.0%) |
| Diploid Tripolar<sup>a</sup> | 1149 (4.7%) | 81 (0.5%) |

<sup>a</sup>To avoid influencing distributions of chromosome counts (Fig. 2), diploid tripolar samples were not required to possess $\geq 10$ aneuploid chromosomes, but are defined by possession $\geq 1$ disomy, $\geq 1$ paternal monosomy, $\geq 1$ maternal monosomy, and $\geq 1$ nullisomy.

Among the different categories of karyotype-wide aberrations, digynic triploid and near-triploid patterns were relatively prevalent (1061 samples or 2.5%; Table 2). Triploid zygotes are predisposed to form tripolar spindles (23), in which case the resulting embryos may be expected to be approximately or ‘quasi-’ diploid with chaotic chromosome complements. This follows from a model by which the triploid set of chromosomes (23 × 3 = 69) is replicated (69 × 2 = 138), then randomly segregated to three daughter cells (138 / 3 = 46 chromosomes per cell). Samples exhibiting this pattern (see Methods) were rare in the PGT-A dataset (106 samples or 0.25%), potentially indicating that chromosome segregation in triploid cells is not generally random.

### Evidence of frequent tripolar mitosis in cleavage-stage embryos

Meanwhile, random tripolar division of normally-fertilized dipronuclear (2PN) zygotes would be predicted to generate hypodiploid complements in the three daughter cells (92 / 3 = ∼31 chromosomes per cell). Under this model, one would expect to observe a mixture of disomies, paternal monosomies, maternal monosomies, and nullisomies in the ratio of 4:2:2:1 (Fig. 2A). Selecting aneuploid samples with at least one disomy, paternal monosomy, maternal monosomy, and nullisomy (and no-co-occurring trisomy or uniparental disomy), we identified 1230 (2.9%) putative ‘diploid tripolar’ samples with a mean number of 28.0 ± 6.3 (±SD) total chromosomes (mode = 30 chromosomes; Fig. 2B). Performing 10,000 simulations, we confirmed that the observed patterns of chromosome loss were consistent with a model of tripolar segregation of diploid cells (Fig. 2C). We note that the distribution of total chromosomes exhibits modest bimodality, with a secondary peak at 20 chromosomes (Fig. 2B). This may reflect an additional round of tripolar mitosis, whereby the diploid tripolar complement is replicated (30.67 × 2 = 61.3) then randomly divided among three daughter cells (61.3 / 3 = 20.4 chromosomes per cell). The high degree of concordance between the observed and simulated data indicates that during tripolar mitosis, chromosomes tend to assort randomly into each of the three daughter cells rather than segregating preferentially to one or two cells. Consistent with the notion that multipolar mitotic divisions are incompatible with development beyond the cleavage stage, more than 90% (1149 samples) of the 1230 putative diploid tripolar samples were from day-3 blastomere biopsies, with only a small minority (81 samples) represented among the trophectoderm biopsies.

**Figure 2.**
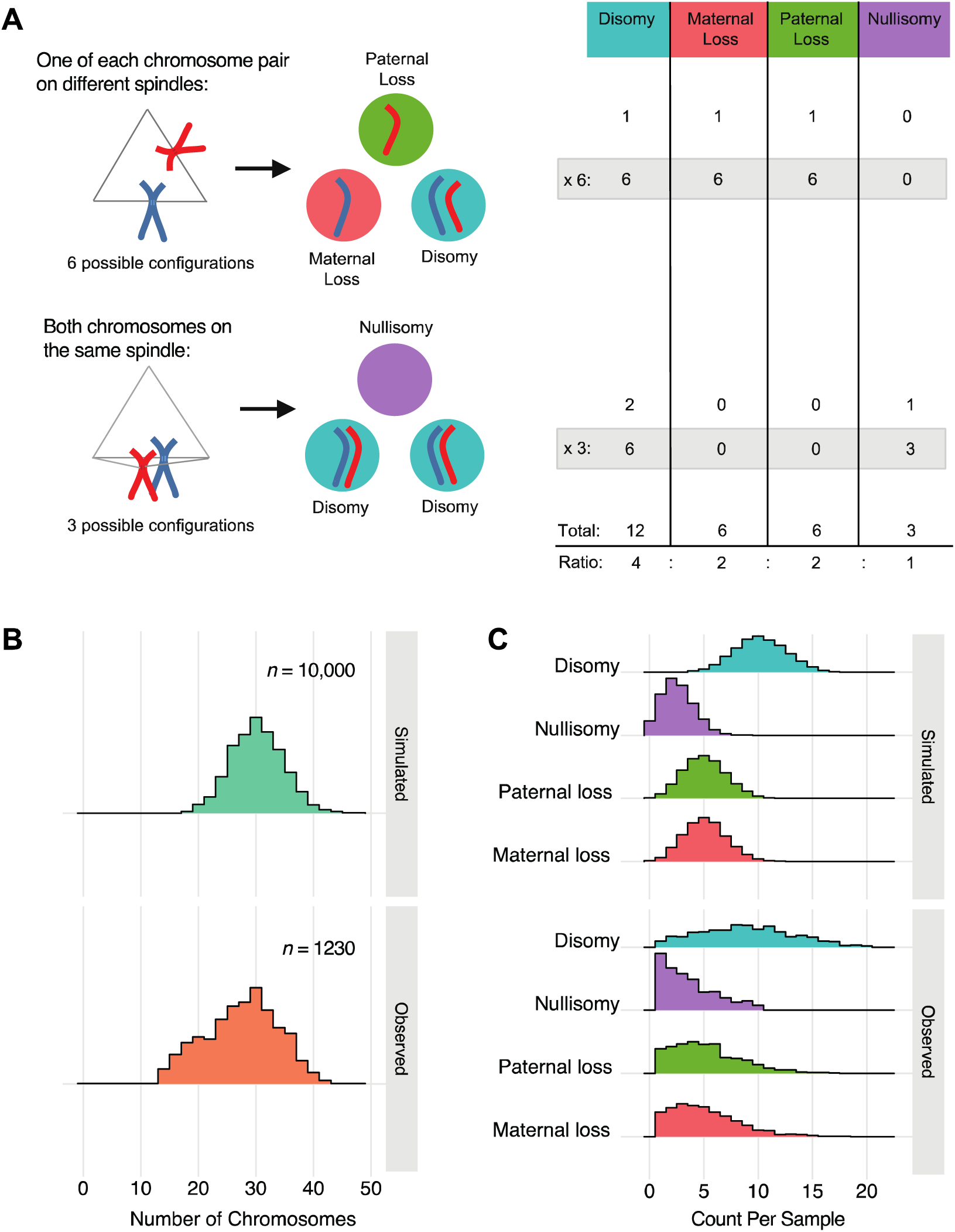
Tripolar mitosis in diploid embryos. (A) Schematic showing all possible outcomes following diploid tripolar mitosis. Only one pair of chromosomes is depicted for clarity (maternal: red; paternal: blue). Spindle attachment configurations are shown on the left with segregation outcomes in the three daughter cells shown in the middle. The table on the right provides the expected ratios of segregation outcomes for each chromosome following tripolar mitosis. (B) Density histograms showing observed chromosome counts in 1230 samples that are suspected to have undergone tripolar mitosis (see main text) compared to 10,000 samples simulated under a model of random tripolar segregation. Observed peaks at ∼30 and ∼20 chromosomes may reflect one and two rounds of tripolar mitosis, respectively. (C) Density histograms displaying the number of chromosomes in each biopsy displaying each of the four possible outcomes. Simulated data (10,000 simulations) are displayed in the top panel, while observed data (1230 putative diploid tripolar biopsies) are displayed in the bottom panel.

### Common maternal variants spanning PLK4 are associated with tripolar mitosis

We recently reported that a common (∼30% global minor allele frequency) haplotype spanning *PLK4*, tagged by the SNP rs2305957, is associated with mitotic error in human preimplantation embryos (28). In addition to the overall association with mitotic-origin aneuploidy, we noted a particularly strong association with complex errors involving loss of both maternal and paternal homologs (28). As the diploid tripolar signature observed in our current analysis resembled this pattern, we sought to test whether the maternal genotype association at *PLK4* was driven by tripolar mitosis of diploid cells.

Supporting our hypothesis, the per-case frequency of diploid tripolar day-3 blastomere samples was positively associated with the minor (A) allele of rs2305957 (quasi-binomial GLM: *OR* = 1.42 [95% CI: 1.29 -1.57], *P* = 1.9 × 10^-12^) (Fig. 3). The magnitude of this association with tripolar mitosis thus exceeds that of the original association with mitotic error (quasi-binomial GLM: *OR* = 1.24 [95% CI: 1.18—1.31], *P* = 6.0 × 10^-15^), despite comprising only approximately one-fifth of all putative mitotic aneuploidies in these samples (1149 vs. 5438). Like the originally described association (28), the effect was additive, with means of 3.5%, 4.7%, and 7.5% diploid tripolar embryos per patient carrying zero, one, and two copies of the risk allele, respectively (Fig. 3). The effect was also constant with maternal age (quasi-binomial GLM: *OR* = 0.99, [95% CI: 0.98 -1.01]; Fig. 3C), consistent with previous results (28). Because the distributions of diploid tripolar samples per case exhibited significant inflation of zeros (AIC-corrected Vuong test: *Z* = −6.61, *P* = 1.9 × 10^−11^), we also fit a hurdle model to the data to account separately for the zero and non-zero portions of the distribution. The hurdle model supported the association both for the zero (binomial GLM: *OR* = 1.41, [95% CI: 1.35 -1.47], *P* < 1 × 10^-10^) and non-zero counts (Poisson GLM: *OR* = 1.31, [95% CI: 1.26 – 1.37], *P* < 1 × 10^−10^).

**Figure 3.**
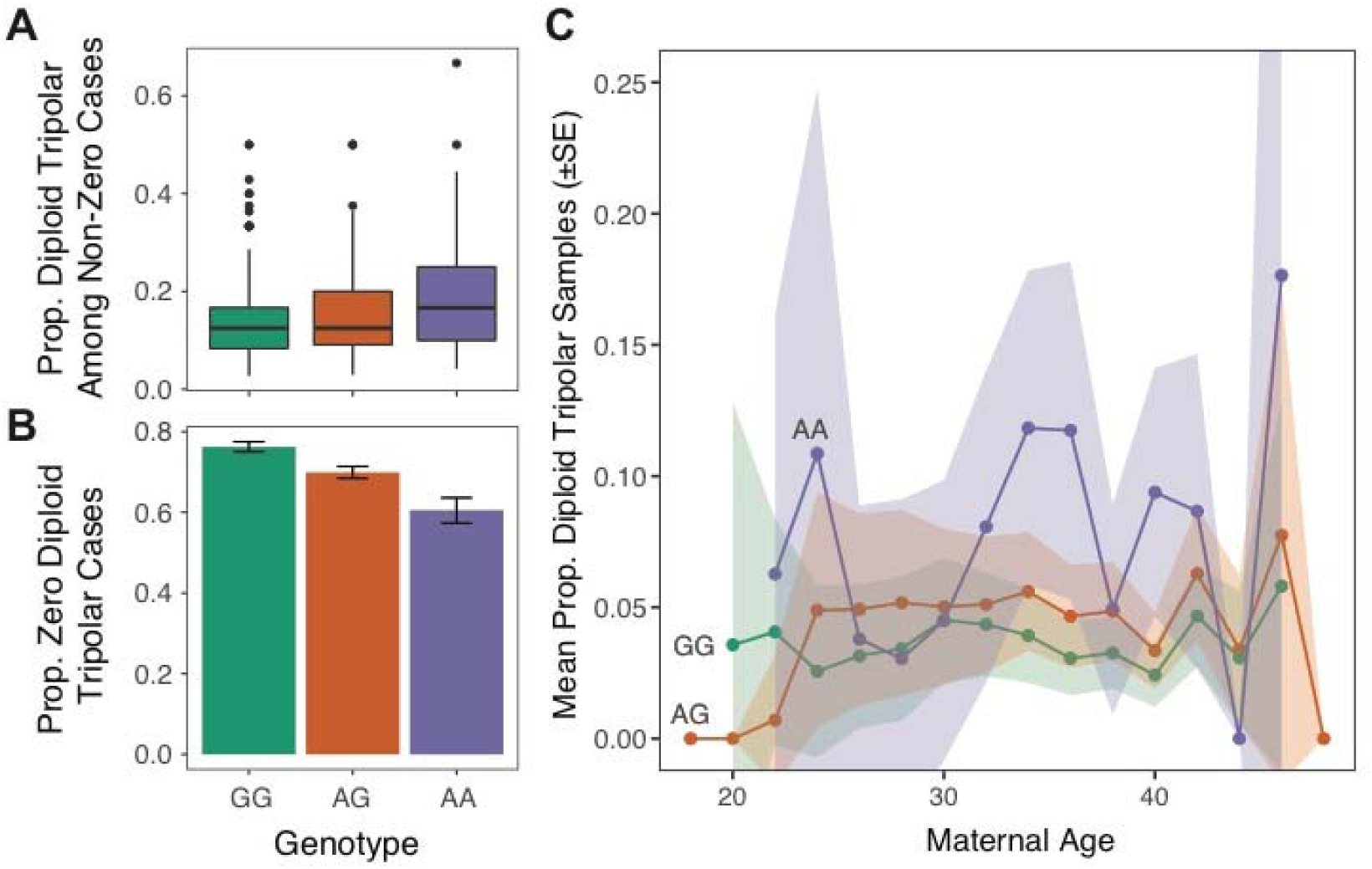
Tripolar mitosis in diploid embryos drives association with common maternal genetic variants spanning *PLK4*. (A) For cases with ≥1 diploid tripolar sample, boxplots of proportion of diploid tripolar samples stratified by maternal genotype at rs2305957. The colored boxes extend from the median to the first (Q1) and third (Q3) quartiles. Whiskers extend to 1.5 * interquartile range. Outlier data points beyond this range are plotted individually. Only cases with >2 samples are plotted for clarity. The risk allele (A allele of rs2305957) is associated with increased frequencies of biopsies exhibiting signatures of tripolar mitosis (quasi-binomial GLM: *OR* = 1.42 [95% CI: 1.29 -1.57], *P* = 1.9 × 10^-12^). (B) Proportions of cases per genotype with zero diploid tripolar samples. Error bars indicate ± standard errors of the mean proportions. (C) Mean proportions of putative diploid tripolar samples stratified by age (rounded to nearest two years) and genotype. Ribbons indicate ± standard errors of the mean proportions.

Upon excluding all putative diploid tripolar samples from the analysis, we re-tested the remaining putative mitotic errors for association with rs2305957 genotype. We observed a modest, but significant association with residual mitotic aneuploidies (quasi-binomial GLM: *OR* = 1.12, [95% CI: 1.06 – 1.19], *P* = 1.6 × 10^-4^), potentially reflecting our initially strict classification of diploid tripolar samples, which excluded co-occurring trisomies. Supporting this hypothesis, significant associations were observed upon excluding the originally defined set of diploid tripolar samples, but relaxing the criteria to include hypodiploid complements (<42 chromosomes) with co-occurring maternal trisomy (quasi-binomial GLM: *OR* = 1.16, [95% CI: 1.04 – 1.30], *P* = 9.5 × 10^-3^) or paternal trisomy (quasi-binomial GLM: *OR* = 1.29, [95% CI: 1.09 – 1.53], *P* = 3.6 × 10^-3^). These results suggest that tripolar mitosis may also occur in embryos already affected by meiotic or mitotic errors or that additional aneuploidies may accumulate downstream of tripolar mitosis.

### Analysis of time-lapse data from a subset of cases

Seeking to validate our findings, we took advantage of the fact that embryos from a subset of IVF cases included in the PGT-A dataset had previously undergone time-lapse imaging by the referring laboratory, with published data demonstrating frequent multipolar mitosis [27] (Supplementary Material, Videos S1, S2, and S3). These included 77 day-3 cleavage-stage embryos from 10 IVF cases with single blastomeres analyzed by PGT-A. Consistent with our hypothesis, we observed that the diploid tripolar PGT-A signature was significantly enriched among embryos documented by time-lapse to have undergone one or more tripolar mitoses during the first three cleavage divisions (Fisher’s Exact Test: *OR* = 6.64, [95% CI: 1.34 – 37.4], *P* = 0.0087). The signature was most prevalent among embryos undergoing tripolar cleavage during the first division (DUC-1) or undergoing multiple tripolar cleavages (DUC-Plus; Fig. 4A), indicating that these cleavage phenotypes may confer a wider distribution of abnormal cells at the time of biopsy.

**Figure 4.**
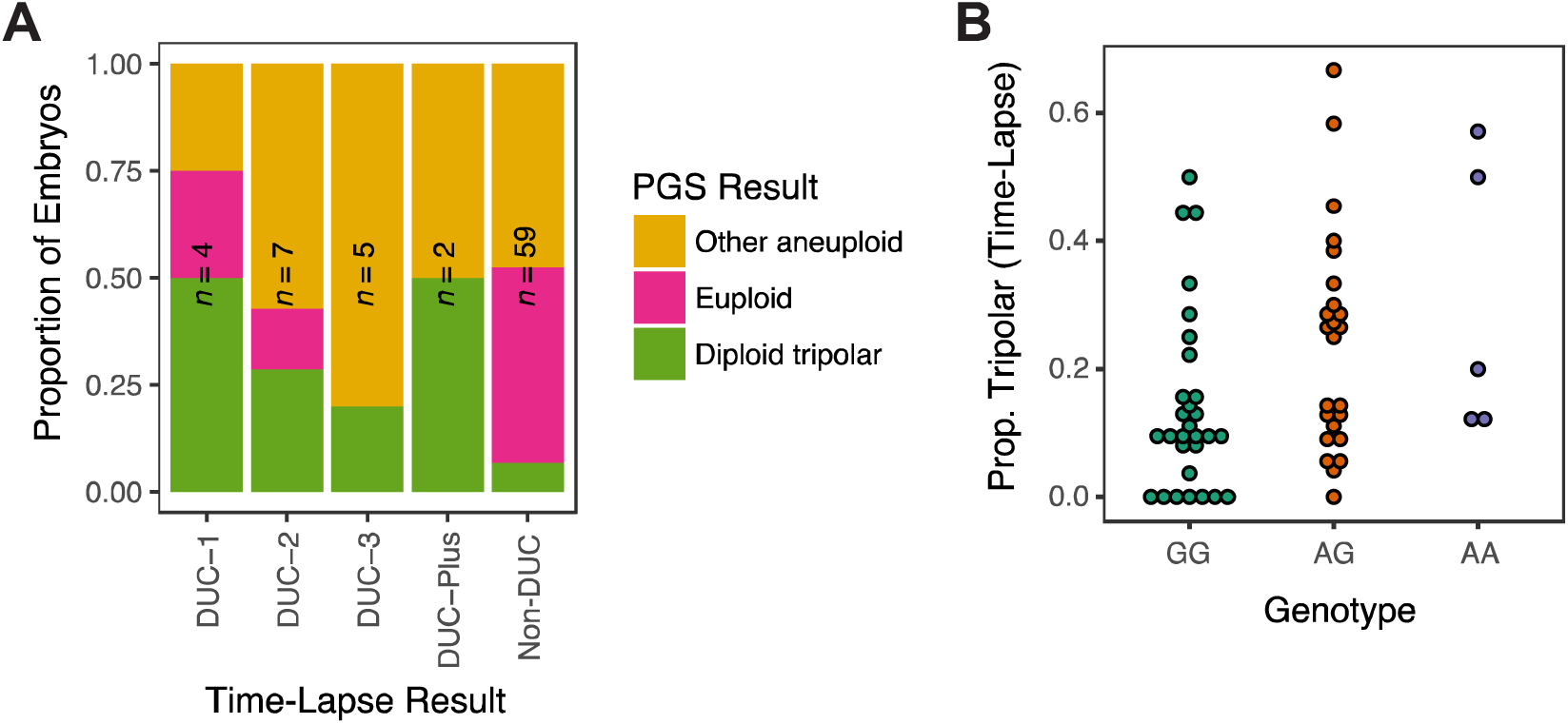
Time-lapse data from a subset of embryos support the PGT-A-based hypothesis of tripolar mitosis and association with *PLK4* genotype. (A) Samples recorded to have undergone tripolar mitosis during the first (DUC-1; Supplementary Material, Video S1), second (DUC-2; Supplementary Material, Video S2), third (DUC-3; Supplementary Material, Video S3) or multiple cleavage divisions are enriched for the diploid tripolar PGT-A signature compared to embryos that underwent normal cleavage (Non-DUC) (Fisher’s Exact Test: *OR* = 6.64, [95% CI: 1.34 – 37.4], *P* = 0.0087). (B) Genotype data from 58 patients whose embryos underwent time-lapse imaging as well as PGT-A analysis of either day-3 or day-5 embryo biopsies support a positive association between the minor allele of rs2305957 and incidence of tripolar mitosis (quasi-binomial GLM: *OR* = 1.53 [95% CI: 1.01 – 2.31], *P* = 0.047).

Time-lapse data were also valuable for direct validation of the association with maternal *PLK4* genotype. For this purpose, data from cases undergoing day-5 blastocyst biopsy could also be incorporated, as the availability of time-lapse data from these cases does not depend on embryo survival to the blastocyst stage. This extended the analysis to a total of 58 IVF patients, with a total of 742 embryos having undergone time-lapse screening [27]. Despite this small sample size, the minor allele of rs2305957 was significantly associated with time-lapse-based incidence of tripolar mitosis (quasi-binomial GLM: *OR* = 1.53 [95% CI: 1.01 – 2.31], *P* = 0.047), with directionality consistent with previous results (Fig. 4B).

## Discussion

Aneuploidy is the leading known cause of implantation failure, miscarriage, and congenital birth defects. Recent studies have vastly improved our understanding of aneuploidy of maternal meiotic origin and its association with maternal age. Meanwhile, relatively little is known about errors of mitotic origin that occur after fertilization and contribute to widespread chromosomal mosaicism. Establishing the molecular mechanisms and risk factors contributing to mitotic aneuploidy may aid the development of future infertility treatments as well as improve our understanding of natural human fertility.

One prominent form of mitotic aneuploidy, termed ‘chaotic mosaicism’, has been recognized since early applications of PGT-A (17), but its origins have proven elusive. Chaotic embryos are characterized by severely aneuploid karyotypes that vary from cell to cell in a pattern reminiscent of cancer cell lines. Intriguingly, the first description of this phenomenon by Delhanty et al. noted that “the occurrence of chaotically dividing embryos was strongly patient-related, i.e. some patients had ‘chaotic’ embryos in repeated cycles, whereas other patients were completely free of this type of anomaly” (17). This observation hints at a patient-specific predisposition to the formation of chaotic embryos, either due to genetic or environmental factors. Technical limitations of the fluorescence *in situ* hybridization (FISH) platform, however, hindered the ability of early studies to examine chaotic embryos in greater detail or probe their molecular origins. Here we revisited this topic equipped with a high-resolution dataset comprising 24-chromosome PGT-A data from >41,000 embryos.

One hypothesized source of chaotic aneuploidy is the formation of multipolar mitotic spindles, a phenomenon frequently observed among triploid embryos. We found that digynic triploidy was common relative to other forms of karyotype-wide abnormality, affecting 3.0% and 1.9% of day-3 and day-5 embryos, respectively. This is consistent with previous studies showing that digynic 3PN zygotes are relatively prevalent and occasionally undetected following ICSI (30), which was used in 80-90% of cases in this dataset. Digynic triploidy may arise either due to failed meiotic division or failed extrusion of the second polar body (26). While previous studies have demonstrated that triploid embryos are predisposed to multipolar cell division, we identified few samples (106 samples or 0.2%) exhibiting patterns suggestive of random chromosome segregation of a replicated digynic triploid genome. This paucity of random triploid tripolar samples agrees with previous reports that digynic 3PN ICSI-derived embryos tend to either remain triploid or self-correct to diploidy (31,32).

Even more prevalent at the cleavage stage (1162 samples or 4.7%) were samples exhibiting hypodiploid karyotypes involving loss of both maternal and paternal homologs. Based on simulation, we showed that this pattern is consistent with a model of random tripolar division of a normal diploid complement, with a subset of cells potentially undergoing a second tripolar division. Time-lapse data from an intersecting set of 77 embryos (27) supported our inference of tripolar mitosis, with the hypodiploid PGT-A signature enriched among embryos previously recorded to have undergone one or more tripolar division. Despite this enrichment, we note that the time-lapse data were not perfectly predictive of the diploid tripolar signature. This result is expected given the sampling noise associated with single-cell biopsy of mosaic embryos. Earlier tripolar divisions would be expected confer a wider distribution of aneuploid cells. Somewhat counterintuitively, however, even one embryo recorded as undergoing tripolar mitosis during the first cleavage (DUC-1) produced a euploid biopsy. Assuming the accuracy of time-lapse and PGT-A classification, this indicates that as an alternative to random segregation, embryos undergoing tripolar mitosis may occasionally segregate normal diploid complement to at least one daughter cell. Interestingly, such a phenomenon was recently documented in a mosaic bovine embryo composed of separate androgenetic, gynogenetic, and normal diploid cell lines (33). The authors proposed that this pattern may have arisen via the formation of a tripolar gonomeric spindle, with separate microtubules associated with maternal and paternal genomes (33).

Notably, we found that diploid tripolar samples were rare at the day-5 blastocyst stage (80 samples or 0.4%). This observation suggests that embryos affected by tripolar mitosis experience reduced viability and/or that aneuploid cell lineages are purged or fail to propagate and thus contribute fewer descendant cells to the blastocyst cell population (34). The former interpretation is consistent with previous studies demonstrating that embryos with complex aneuploidies, including those deriving from normally fertilized zygotes, tend to arrest at cleavage stages, around the time of embryonic genome activation (8,35,36,37). Moreover, previous studies have observed a strong enrichment of abnormal mitotic spindles among arrested cleavage-stage embryos (18), as well as demonstrating that zygotes undergoing tripolar mitosis have poor developmental potential beyond the cleavage stage (24,25,26,27). This hypothesis was recently validated based on combined karyomapping and time-lapse imaging of 25 arrested embryos as well as cells extruded during blastocyst formation of an additional 7 embryos (19). While there is also evidence of tripolar spindle formation in confocal images of fixed human blastocysts (Supplementary Material, Fig. S1), it is possible that aneuploid daughter cells are eliminated by apoptosis or that the spindle configurations are transient and do not tend to result in tripolar anaphase at this stage. Potentially relevant are previous observations that unlike cleavage-stage embryos, adult somatic cells possessing extra centrosomes rarely undergo multipolar mitosis, but instead experience centrosome clustering leading to the formation of a pseudo-bipolar spindle (38). This may reflect increased stringency of mitotic checkpoints following embryonic genome activation, possibly via upregulation of *MAD2* (39). While clustered centrosomes predispose the cell to aberrant microtubule-kinetochore attachments and anaphase lag (40), the resultant aneuploidies are less severe than those induced by multipolar mitosis.

We recently reported that common maternal genetic variants defining a ∼600 Kb haplotype of chromosome 4, tagged by SNP rs2305957, is strongly associated with mitotic aneuploidy in day-3 cleavage-stage embryos. Furthermore, individuals carrying the risk allele had fewer blastocyst-stage embryos available for testing at day 5, suggesting that the aneuploid phenotype impairs embryonic survival. The association we observed was significant with maternal, but not paternal genotypes, reflecting the fact that prior to embryonic genome activation, mitotic divisions are controlled by maternal gene products deposited in the oocyte (28). While the associated haplotype spans seven genes (*INTU*, *SLC25A31*, *HSPA4L*, *PLK4*, *MFSD8*, *LARP1B*, and *PGRMC2*), *PLK4* stands out given its well-characterized role as an essential master regulator of centrosome duplication. *PLK4* is a tightly regulated kinase that initiates assembly of a daughter procentriole at the base of the existing mother centriole, thereby mediating bipolar spindle formation (41,42). Altered expression of *PLK4* is associated with centrosome amplification, generating multipolar spindles in human cell lines—a hallmark of several cancers (41,42,43,44,45). Based on this knowledge, we hypothesized that the tripolar mitosis mechanism operating in cleavage-stage embryos may drive the association between aneuploidy and *PLK4* genotype. Consistent with this hypothesis, the diploid tripolar signature was strongly associated with maternal genotype at rs2305957—a finding that we further validated using time-lapse data from 742 embryos and corresponding genotype data from 58 patients. Together, our results support a causal role of *PLK4* in the original association.

Zhang et al. (46) recently replicated a key finding of our initial association study, demonstrating that rates of blastocyst formation are reduced among embryos from patients carrying the high-risk genotypes of rs2305957. Furthermore, patients diagnosed with early recurrent miscarriage were found to possess a higher frequency of the risk allele compared to matched fertile control subjects (46). Our analysis provides evidence of the molecular mechanism underlying these results, suggesting that complex aneuploidies arising from tripolar mitosis are associated with increased rates of embryonic mortality prior to blastocyst formation. Along with methodological considerations, this may explain the lack of association with recurrent (post-implantation) pregnancy losses (47) and other clinical diagnoses (8). Further research will be necessary to determine the relevance of these findings to non-IVF patient populations and *in vivo* conception.

Through detailed characterization of karyotype-wide abnormalities, our analysis revealed that tripolar mitosis in embryos originating from normally-fertilized 2PN zygotes is an under-recognized phenomenon contributing to aneuploidy in cleavage-stage embryos. Using simulation, we showed that observed chromosomal patterns are consistent with a model of random segregation of a replicated diploid complement to each of three daughter cells. Finally, we demonstrated that the incidence of these tripolar mitoses is patient-specific and significantly correlated with maternal genetic variants spanning the centrosomal regulator *PLK4*. This implicates maternal variation in *PLK4* as a factor influencing mitotic spindle integrity while shedding light on tripolar mitosis as an important mechanism contributing to aneuploidy in preimplantation embryos. Together our results help illuminate the molecular origins of chaotic aneuploidy, a long-recognized phenomenon contributing to high rates of IVF failure.

## Materials and Methods

### Human subjects approvals

Based on the retrospective nature of the analysis and use of de-identified data, this work was determined not to constitute human subjects research by the University of Washington Human Subjects Division as well as Ethical & Independent Review Services, who provided their determination to Natera, Inc.

### Sample preparation, genotyping, and aneuploidy detection

DNA isolation, whole genome amplification, and SNP genotyping are described in detail in McCoy et al. (8). Briefly, genetic material was obtained from IVF patients or oocyte donors and male partners by buccal swab or peripheral venipuncture. Genetic material was also obtained from single blastomere biopsies of day-3 cleavage-stage embryos or ∼5-10 cell biopsies of trophectoderm tissue from day-5 blastocysts. Embryo DNA was amplified via multiple displacement amplification (MDA; see [8]). Amplified embryo DNA and bulk parental tissue were genotyped on the HumanCytoSNP-12 BeadChip (Illumina; San Diego, CA) using the standard Infinium II protocol. Genotype calling was performed using the GenomeStudio software package (Illumina; San Diego, CA).

Aneuploidy detection was performed using the Parental Support algorithm, which leverages informative parental markers (e.g. sites where one parent is homozygous and the other parent is heterozygous) to infer transmission of maternal and paternal homologs along with the copy number of all chromosomes across the embryo genome. In doing so, the method overcomes high rates of allelic dropout that characterize data obtained from whole genome amplified DNA of embryo biopsies. The Parental Support method is described in detail in Johnson et al. (9), who also demonstrated its sensitivity and specificity of 97.9% and 96.1%, respectively.

### Classification criteria

Putative diploid tripolar samples were identified as those exhibiting ≥1 maternal monosomy, ≥1 paternal monosomy, ≥1 nullisomy, and ≥1 disomy with no co-occurring trisomies or uniparental disomies. Other karyotype-wide abnormalities (Table 2) were required to exhibit ≥10 aneuploid chromosomes. Digynic and diandric triploid and near-triploid samples were defined as those with ≥20 maternal trisomic or paternal trisomic chromosomes, respectively. Maternal haploid and paternal haploid or near-haploid samples were similarly defined as those with ≥20 paternal monosomic or maternal monosomic chromosomes, respectively. Digynic triploid tripolar and diandric triploid tripolar samples were defined using simulation-based expectations of random chromosome segregation of triploid cells. Specifically, digynic triploid tripolar samples were required to exhibit ≥3 disomic, ≥3 maternal trisomic, 1-8 maternal uniparental disomic, and 1-8 paternal monosomic chromosomes. Diandric triploid tripolar samples were meanwhile required to exhibit ≥3 disomic, ≥3 paternal trisomic, 1-8 paternal uniparental disomic, and 1-8 maternal monosomic chromosomes.

### Time-lapse imaging and annotation

Time-lapse microscope image capture, annotation, and classification of tripolar mitosis (therein referred to as ‘direct unequal cleavage’ or ‘DUC’), is described in the Materials and Methods section of Zhan et al. (27). In short, images were automatically captured every 10 minutes, with seven focal planes illuminated by 635 nm LED light. Several time points were annotated, including: appearance of pronuclei, syngamy, time of division, morula, cavitation, early blastocyst, expanded blastocyst, and hatching blastocyst. Only cells with visible nuclei were considered blastomeres. DUC was defined as 1) cleavage of a single blastomere into 3+ daughter blastomeres or 2) unusually short interval between mother and daughter cell division of ≤5 hours, in accordance with established criteria [48,49]. DUC-1 (first cleavage; Supplementary Material, Video S1), DUC-2 (second cleavage; Supplementary Material, Video S2), DUC-3 (third cleavage; Supplementary Material, Video S3), and DUC-Plus (multiple DUCs) were annotated based on the cleavage stage during which DUC was observed.

### Statistical analyses

Statistical analyses were performed using the R statistical computing environment (50), with plots generated using the ‘ggplot2’ package (51). Density plots (Fig. 2A and B) were generated using the ggjoy package (https://cran.r-project.org/package=ggjoy). To test the association between maternal genotype and incidence of embryos with the diploid tripolar PGT-A signature, we limited the analysis to the originally tested set of unrelated IVF patients or egg donors (no repeat cases; less than second degree relatedness) with genotype data meeting quality control thresholds (95% variant call rate; 95% sample genotyping efficiency). We then used a generalized linear regression model to test for association between the per-case counts of samples that did and did not exhibit the diploid tripolar pattern and maternal genotypes at rs2305957, encoded as dosage of the ‘A’ allele. To account for excess zeros, we also fit a hurdle model to the zero and non-zero portions of the distribution using the ‘pscl’ package (52). For the corresponding analysis of time-lapse phenotypes, we sought to maximize statistical power from a small number of cases by combining rather than excluding embryo data from repeat IVF cases. To this end, we used KING (53) to infer cases derived from the same patient based on maternal genotype data. We then summed tripolar and non-tripolar embryo phenotypes for each independent patient, using the resulting data in a quasi-binomial regression, as above. Aneuploidy data were made available with the original publication (28), posted as supplementary materials. Analysis scripts are posted on GitHub https://github.com/rmccoy7541/tripolar_mitosis.

## Acknowledgements

The authors would like to thank Allison Ryan, Milena Banjevic, Matthew Hill, and Matthew Rabinowitz for their work at Natera to develop algorithms to detect aneuploidies in preimplantation genetic screening data. We also thank Molly Gasperini and members of the Akey and Shendure labs for helpful discussions on this project.

## Funding

This work was supported in part by NIH/NHGRI Genome Training Grant (5T32HG000035-22) to the Department of Genome Sciences at the University of Washington.

## References

1. Fragouli, E., Alfarawati, S., Spath, K., Jaroudi, S., Sarasa, J., Enciso, M. and Wells, D. (2013). The origin and impact of embryonic aneuploidy. Hum. Genet., 132, 1001–1013.

2. Hassold, T., Hall, H. and Hunt, P. (2007). The origin of human aneuploidy: where we have been, where we are going. Hum. Mol. Genet., 16, R203–R208.

3. Handyside, A.H., Montag, M., Magli, M.C., Repping, S., Harper, J., Schmutzler, A., Vesela, K., Gianaroli, L. and Geraedts, J. (2012). Multiple meiotic errors caused by predivision of chromatids in women of advanced maternal age undergoing in vitro fertilisation. Eur. J. Hum. Genet., 20, 742–747.

4. Vanneste, E., Voet, T., Le Caignec, C., Ampe, M., Konings, P., Melotte, C., Debrock, S., Amyere, M., Vikkula, M., Schuit, F., et al. (2009). Chromosome instability is common in human cleavage-stage embryos. Nat. Med., 15, 577–583.

5. Handyside, A.H. (2013). 24-chromosome copy number analysis: a comparison of available technologies. Fertil. Steril., 100, 595–602.

6. Zegers-Hochschild, F., Adamson, G.D., Dyer, S., Racowsky, C., de Mouzon, J., Sokol, R., Rienzi, L., Sunde, A., Schmidt, L., Cooke, I.D. et al. (2017). The International Glossary on Infertility and Fertility Care, 2017. Fertil. Steril., in press.

7. Donate, A., Estop, A. M., Giraldo, J. and Templado, C. (2016). Paternal age and numerical chromosome abnormalities in human spermatozoa. Cytogenet. Genome Res., 148, 241–248.

8. McCoy, R.C., Demko, Z.P., Ryan, A., Banjevic, M., Hill, M., Sigurjonsson, S., Rabinowitz, M. and Petrov, D.A. (2015). Evidence of selection against complex mitotic-origin aneuploidy during preimplantation development. PLoS Genet., 11, e1005601.

9. Johnson, D.S., Gemelos, G., Baner, J., Ryan, A., Cinnioglu, C., Banjevic, M., Ross, R., Alper, M., Barrett, B., Frederick, J., et al. (2010). Preclinical validation of a microarray method for full molecular karyotyping of blastomeres in a 24-h protocol. Hum. Reprod., 25, 1066–1075.

10. Rabinowitz, M., Ryan, A., Gemelos, G., Hill, M., Baner, J., Cinnioglu, C., Banjevic, M., Potter, D., Petrov, D.A. and Demko, Z. (2012). Origins and rates of aneuploidy in human blastomeres. Fertil. Steril., 97, 395–401.

11. Handyside, A.H., Harton, G.L., Mariani, B., Thornhill, A.R., Affara, N., Shaw, M.A. and Griffin, D.K. (2010). Karyomapping: a universal method for genome wide analysis of genetic disease based on mapping crossovers between parental haplotypes. J. Med. Genet., 47, 651–658.

12. Ottolini, C.S., Newnham, L.J., Capalbo, A., Natesan, S.A., Joshi, H.A., Cimadomo, D., Griffin, D.K., Sage, K., Summers, M.C., Thornhill, A.R., et al. (2015). Genome-wide maps of recombination and chromosome segregation in human oocytes and embryos show selection for maternal recombination rates. Nature Genet., 47, 727–735.

13. Natesan, S.A., Bladon, A.J., Coskun, S., Qubbaj, W., Prates, R., Munne, S., Coonen, E., Dreesen, J.C., Stevens, S.J., Paulussen, A.D. et al. (2014). Genome-wide karyomapping accurately identifies the inheritance of single-gene defects in human preimplantation embryos in vitro. Genet. Med., 16, 838–845.

14. Kumar, A., Ryan, A., Kitzman, J.O., Wemmer, N., Snyder, M.W., Sigurjonsson, S., Lee, C., Banjevic, M., Zarutskie, P.W., Lewis, A.P. et al. (2015). Whole genome prediction for preimplantation genetic diagnosis. Genome Med., 7, 35.

15. Hou, Y., Fan, W., Yan, L., Li, R., Lian, Y., Huang, J., Li, J., Xu, L., Tang, F., Xie, X.S. et al. (2013). Genome analyses of single human oocytes. Cell, 155, 1492–1506.

16. Mantikou, E., Wong, K.M., Repping, S. and Mastenbroek, S. (2012). Molecular origin of mitotic aneuploidies in preimplantation embryos. Biochim. Biophys. Acta, 1822, 1921–1930.

17. Delhanty, J.D., Harper, J.C., Ao, A., Handyside, A.H. and Winston, R.M. (1997). Multicolour FISH detects frequent chromosomal mosaicism and chaotic division in normal preimplantation embryos from fertile patients. Hum. Genet., 99, 755–760.

18. Chatzimeletiou, K., Morrison, E.E., Prapas, N., Prapas, Y. and Handyside, A.H. (2005). Spindle abnormalities in normally developing and arrested human preimplantation embryos in vitro identified by confocal laser scanning microscopy. Hum. Reprod., 20, 672–682.

19. Ottolini, C., Kitchen, J., Xanthopoulou, L., Gordon, T., Summers, M.C. and Handyside, A.H. (2017). Tripolar mitosis and partitioning of the genome arrests human preimplantation development in vitro. Sci. Rep., 7, 9744.

20. Boveri, T. (1900). Zellen-Studien: Heft 4, Ueber die natur der centrosomen. Verlag Von Gustav Fischer, Stuttgart, Germany.

21. Balczon, R., Bao, L., Zimmer, W.E., Brown, K., Zinkowski, R.P. and Brinkley, B.R. (1995). Dissociation of centrosome replication events from cycles of DNA synthesis and mitotic division in hydroxyurea-arrested Chinese hamster ovary cells. J. Cell. Biol., 130, 105–115.

22. Plachot, M., Mandelbaum, J., Junca, A.M., De Grouchy, J., Salat-Baroux, J. and Cohen, J. (1989). Cytogenetic analysis and developmental capacity of normal and abnormal embryos after IVF. Hum. Reprod., 4, 99–103.

23. Kola, I., Trounson, A., Dawson, G. and Rogers, P. (1987). Tripronuclear human oocytes: altered cleavage patterns and subsequent karyotypic analysis of embryos. Biol. Reprod., 37, 395–401.

24. Wirka, K.A., Chen, A.A., Conaghan, J., Ivani, K., Gvakharia, M., Behr, B., Suraj, V., Tan, L. and Shen, S. (2014). Atypical embryo phenotypes identified by time-lapse microscopy: high prevalence and association with embryo development. Fertil. Steril., 101, 1637–1648.

25. Hlinka, D., Kalatova, B., Uhrinova, I., Dolinska, S., Rutarova, J., Rezacova, J., Lazarovska, S. and Dudas, M. (2012). Time-lapse cleavage rating predicts human embryo viability. Physiol. Res., 61, 513.

26. Kalatova, B., Jesenska, R., Hlinka, D. and Dudas, M. (2015). Tripolar mitosis in human cells and embryos: Occurrence, pathophysiology and medical implications. Acta Histochem., 117, 111–125.

27. Zhan, Q., Ye, Z., Clarke, R., Rosenwaks, Z. and Zaninovic, N. (2016). Direct Unequal Cleavages: Embryo Developmental Competence, Genetic Constitution and Clinical Outcome. PLoS One, 11, p.e0166398.

28. McCoy, R.C., Demko, Z., Ryan, A., Banjevic, M., Hill, M., Sigurjonsson, S., Rabinowitz, M., Fraser, H.B. and Petrov, D.A. (2015). Common variants spanning PLK4 are associated with mitotic-origin aneuploidy in human embryos. Science, 348, 235–238.

29. McCoy, R.C. (2017). Mosaicism in Preimplantation Human Embryos: When Chromosomal Abnormalities Are the Norm. Trends Genet., 33, 448–463.

30. Staessen, C. and Van Steirteghem, A.C. (1997). The chromosomal constitution of embryos developing from abnormally fertilized oocytes after intracytoplasmic sperm injection and conventional in-vitro fertilization. Hum. Reprod., 12, 321–327.

31. Grau, N., Escrich, L., Martín, J., Rubio, C., Pellicer, A. and Escribá, M.J. (2011). Self-correction in tripronucleated human embryos. Fertil. Steril., 96, 951–956.

32. Grau, N., Escrich, L., Galiana, Y., Meseguer, M., García-Herrero, S., Remohí, J., and Escribá, M. J. (2015). Morphokinetics as a predictor of self-correction to diploidy in tripronucleated intracytoplasmic sperm injection–derived human embryos. Fertil. Steril., 104, 728–735.

33. Destouni, A., Esteki, M.Z., Catteeuw, M., Tšuiko, O., Dimitriadou, E., Smits, K., Kurg, A., Salumets, A., Van Soom, A., Voet, T. et al. (2016). Zygotes segregate entire parental genomes in distinct blastomere lineages causing cleavage-stage chimerism and mixoploidy. Genome Res., 26, 567–578.

34. Bolton, H., Graham, S.J., Van der Aa, N., Kumar, P., Theunis, K., Gallardo, E. F., Voet, T. and Zernicka-Goetz, M. (2016). Mouse model of chromosome mosaicism reveals lineage-specific depletion of aneuploid cells and normal developmental potential. Nat. Comm., 7, 11165.

35. Sandalinas, M., Sadowy, S., Alikani, M., Calderon, G., Cohen, J. and Munné, S. (2001). Developmental ability of chromosomally abnormal human embryos to develop to the blastocyst stage. Hum. Reprod., 16, 1954–1958.

36. Ruangvutilert, P., Delhanty, J.D., Serhal, P., Simopoulou, M., Rodeck, C.H. and Harper, J.C. (2000). FISH analysis on day 5 post-insemination of human arrested and blastocyst stage embryos. Prenat. Diagn., 20, 552–560.

37. Rubio, C., Rodrigo, L., Mercader, A., Mateu, E., Buendía, P., Pehlivan, T., Viloria, T., De los Santos, M.J., Simón, C., Remohí, J. and Pellicer, A. (2007). Impact of chromosomal abnormalities on preimplantation embryo development. Prenat. Diagn., 27, 748–756.

38. Quintyne, N.J., Reing, J.E., Hoffelder, D.R., Gollin, S.M. and Saunders, W.S. (2005). Spindle multipolarity is prevented by centrosomal clustering. Science, 307, 127–129.

39. Krämer, A., Maier, B. and Bartek, J. (2011). Centrosome clustering and chromosomal (in) stability: a matter of life and death. Mol. Oncol., 5, 324–335.

40. Ganem, N.J., Godinho, S.A. and Pellman, D. (2009). A mechanism linking extra centrosomes to chromosomal instability. Nature, 460, 278–282.

41. Habedanck, R., Stierhof, Y.D., Wilkinson, C.J. and Nigg, E.A. (2005). The Polo kinase Plk4 functions in centriole duplication. Nat. Cell Biol., 7, 1140–1146.

42. Bettencourt-Dias, M., Rodrigues-Martins, A., Carpenter, L., Riparbelli, M., Lehmann, L., Gatt, M.K., Carmo, N., Balloux, F., Callaini, G. and Glover, D.M. (2005). SAK/PLK4 is required for centriole duplication and flagella development. Current Biol., 15, 2199–2207.

43. Ko, M.A., Rosario, C.O., Hudson, J.W., Kulkarni, S., Pollett, A., Dennis, J.W. and Swallow, C.J. (2005). Plk4 haploinsufficiency causes mitotic infidelity and carcinogenesis. Nature Genet., 37, 883.

44. Duensing, A., Liu, Y., Perdreau, S.A., Kleylein-Sohn, J., Nigg, E.A. and Duensing, S. (2007). Centriole overduplication through the concurrent formation of multiple daughter centrioles at single maternal templates. Oncogene, 26, 6280–6288.

45. Rosario, C.O., Ko, M.A., Haffani, Y.Z., Gladdy, R.A., Paderova, J., Pollett, A., Squire, J.A., Dennis, J.W. and Swallow, C.J. (2010). Plk4 is required for cytokinesis and maintenance of chromosomal stability. Proc. Natl. Acad. Sci. USA, 107, 6888–6893.

46. Zhang, Q., Li, G., Zhang, L., Sun, X., Zhang, D., Lu, J., Ma, J., Yan, J. and Chen, Z.J. (2017). Maternal common variant rs2305957 spanning PLK4 is associated with blastocyst formation and early recurrent miscarriage. Fertil. Steril., 107, 1034–1040.

47. Sharif, F.A. and Ashour, M. (2015). The single nucleotide polymorphism rs2305957 G/A is not associated with recurrent pregnancy loss. Int. J. Res. Med. Sci., 3, 3123–3125.

48. Meseguer, M., Herrero, J., Tejera, A., Hilligsøe, K.M., Ramsing, N.B. and Remohí, J. (2011). The use of morphokinetics as a predictor of embryo implantation. Hum. Reprod., 26, 2658–2671.

49. Rubio, I., Kuhlmann, R., Agerholm, I., Kirk, J., Herrero, J., Escribá, M.J., Bellver, J. and Meseguer, M. (2012). Limited implantation success of direct-cleaved human zygotes: a time-lapse study. Fertil. Steril., 98, 1458–1463.

50. R Development Core Team. (2013). R: A Language and Environment for Statistical Computing. R Foundation for Statistical Computing, Vienna, Austria.

51. Wickham, H. (2016). ggplot2: elegant graphics for data analysis. Springer, New York, NY.

52. Zeileis, A., Kleiber, C., Jackman, S. (2008). Regression models for count data in R. J. Stat. Softw., 27, 1–25.

53. Manichaikul, A., Mychaleckyj, J. C., Rich, S. S., Daly, K., Sale, M., Chen, W. M. (2010). Robust relationship inference in genome-wide association studies. Bioinformatics, 26, 2867–2873.

